# Lysyl oxidase-mediated intermolecular crosslinks fine-tune collagen I molecular dynamics and regulate cell-matrix interactions through focal adhesions

**DOI:** 10.1101/2025.08.25.672158

**Authors:** Scott Dillon, Jonathan Clark, Melinda J. Duer

## Abstract

Lysyl oxidase (LOX)–mediated intermolecular crosslinking is essential for collagen I fibril stability, yet its influence on collagen molecular conformation and dynamics, and the downstream consequences for cell–matrix interactions remain poorly understood. Here, we genetically modulated LOX in collagen I–producing MC3T3-E1 cells to generate matrices with elevated (overexpression, OX) or absent (knockout, KO) crosslinking. Enhanced crosslinking yielded thick, continuous, aligned fibrils, whereas reduced crosslinking produced friable, dissociated fibrils. Solid-state nuclear magnetic resonance spectroscopy (SSNMR) revealed local triple-helix unfolding and altered nanosecond- and microsecond-scale molecular motions in both OX and KO matrices, changes largely reversible upon decellularization, implicating a synergistic role of crosslinking chemistry and cell-applied forces in regulating the dynamically-accessible conformations of collagen I. Changes in the molecular structure and dynamics of collagen had a functional impact on cell adhesion and mechanotransduction. These findings identify collagen crosslinking as a tunable element of the extracellular matrix “mechanical code,” integrating biochemical modification with molecular-scale mechanics to regulate cell-matrix adhesion and mechanosignalling.

## Introduction

Collagen type I is the principal structural protein in most vertebrate connective tissues and plays a central role in maintaining the mechanical integrity of biological tissues. Its unique triple-helical structure assembles into fibrils and larger fibers that provide tensile strength and resistance to mechanical deformation^1–3^. However, the full mechanical performance of collagen-rich tissues is not solely dictated by the collagen molecules themselves, but also by covalent crosslinks that form between them. These crosslinks are largely introduced through the action of lysyl oxidase (LOX), a copper-dependent amine oxidase which catalyzes the oxidative deamination of lysine and hydroxylysine residues in collagen I^4,5^. This enzymatic step initiates the formation of reactive (hydroxy)allysine aldehydes which participate in spontaneous condensation reactions to form covalent crosslinks between adjacent collagen molecules^6^.

The accumulation and maturation of LOX-mediated crosslinks significantly influence the bulk mechanical properties of collagenous tissues. Increased crosslinking enhances tensile strength and stiffness while reducing solubility and enzymatic degradability. These effects are essential for normal development, tissue remodelling, and homeostasis. However, dysregulated crosslinking has been implicated in a variety of pathological conditions. Excessive or abnormal crosslinking is a hallmark of fibrotic diseases, where it contributes to tissue stiffening and functional impairment, and is also commonly observed in stromal tumour and metastatic microenvironments^7–11^. In these contexts, changes in extracellular matrix (ECM) mechanical properties and architecture can profoundly influence cell behaviour by altering cell-matrix interactions, mechanotransduction and cell signaling^9,12,13^.

Despite extensive study of the mechanical and biochemical consequences of LOX-mediated crosslinking at the tissue level, much less is known about its influence on the molecular conformation and dynamic behaviour of individual tropocollagen molecules within fibrils.

Crosslinking may restrict local molecular motion and alter the flexibility of the triple helix^14^. Even subtle changes in collagen molecular geometry as a result of crosslinking can potentially influence higher-order fibrillar organization and affect the mechanical response of the matrix at larger scales^15,16^, because of the highly ordered and periodic arrangement of collagen molecules within fibrils.

These molecular-level alterations may have important implications for cell-matrix interactions. Collagen-binding receptors such as integrins and discoidin domain receptors (DDRs) recognize specific sites on collagen fibrils and initiate intracellular signalling cascades which regulate adhesion, migration, and differentiation^17,18^. If LOX-mediated crosslinking changes the conformational accessibility or dynamics of these binding motifs, it may alter receptor binding kinetics and downstream signalling outcomes. Therefore, understanding how enzymatic crosslinking affects the structural and dynamic properties of collagen at the molecular level is crucial for linking matrix chemistry to cell behaviour and tissue function. However, this aspect of collagen structural biology remains poorly understood.

## Results

### Intermolecular crosslinking alters collagen microarchitecture

To establish the effect of altered LOX expression and consequent collagen crosslink density, we engineered MC3T3-E1 collagen I-producing cells either overexpressing LOX (OX) or harbouring a homozygous frameshift mutation resulting in a premature stop codon in exon 1 of the *Lox* gene rendering the protein non-functional (knock-out; KO). Quantitative reverse transcription polymerase chain reaction (RT-qPCR) in resulting clones verified a significant increase (p < 0.05) in *Lox* expression in OX samples and decrease (p < 0.05) in KO samples (Supp. Fig. 1) compared to wild-type (WT) controls. Immunofluorescence did not show any aberrant localisation of LOX in OX cells and the expected reduction in intensity in KO cells (Supp. Fig. 1).

To establish the effect of altered crosslinking on the microarchitecture of collagen I, cells were stimulated to generate *in vitro* matrices with L-ascorbic acid over 21 days of culture and collagen organisation visualised using immunofluorescence. Compared to WT samples, collagen fibres in OX matrices were dense and more oriented, while KO fibres appeared friable with fewer large fibres (Fig. 1, A). Scanning electron microscopy (SEM) in decellularized samples was used to investigate the matrix nanostructure, and revealed thickened fibres comprised of many tightly bound individual fibrils in parallel in OX samples (Fig. 1, B; white arrowheads), compared to KO samples where thin fibrils exhibited less frequent association with one another (Fig. 1, B).

**Figure 1:**
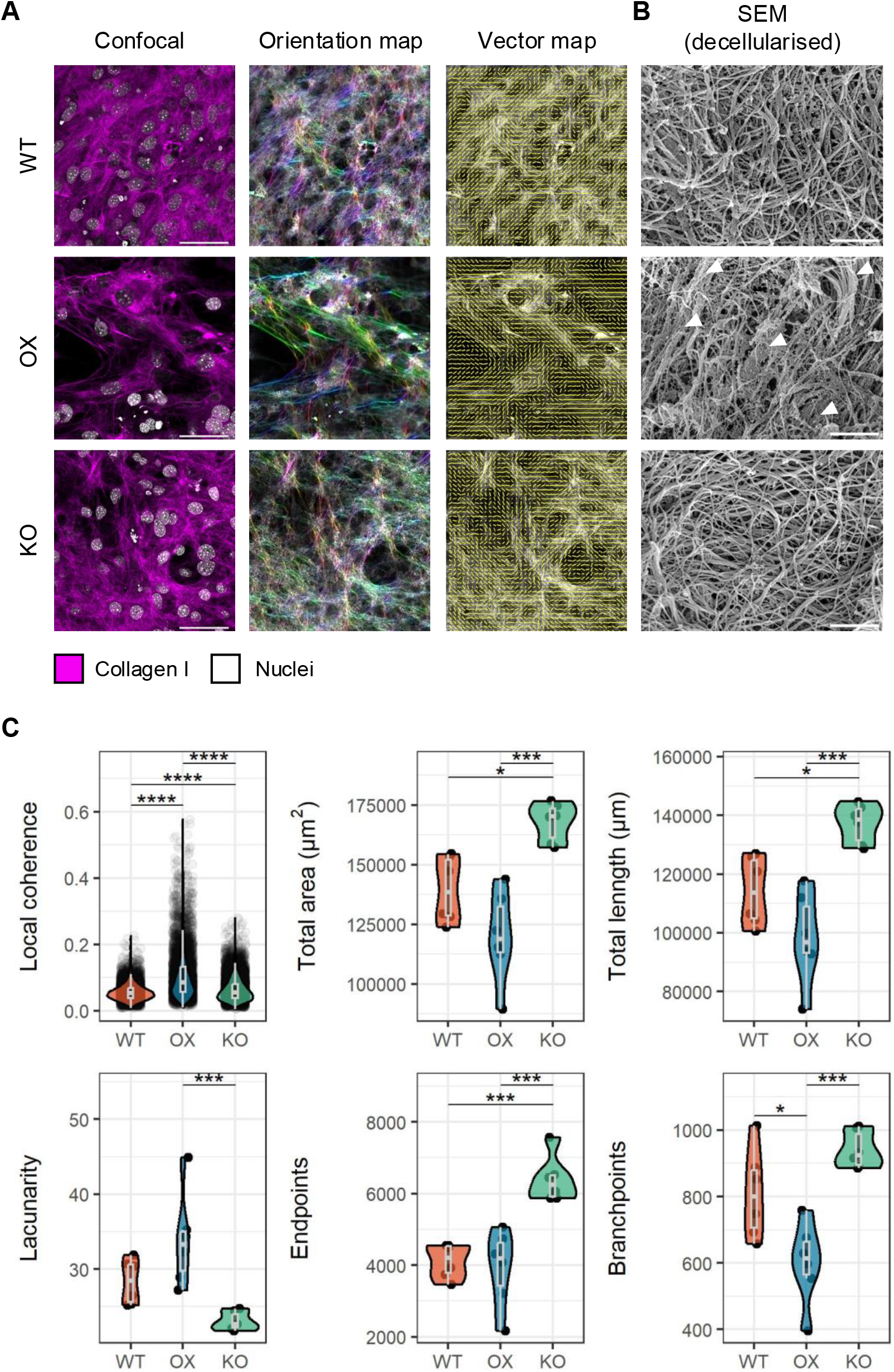
Expression of lysyl oxidase induces changes in collagen I microarchitecture. **(A)** Confocal microscopy for collagen I in edited MC3T3 osteoblasts and local orientation analysis using OrientationJ. **(B)** Scanning electron microscopy (SEM) in decellularised *in vitro* matrix samples. **(C)** Quantification of local coherence and collagen fibre parameters using confocal images. Scales bars = 40µm (A); 1µm (B). In imaging data n = 6 images/genotype from at least 2 independent experiments. Statistical testing performed using pairwise Tukey Honest Significant Difference tests with Bonferonni correction: ^*^ p < 0.05; ^**^ p < 0.01; ^***^ p < 0.001; ^****^ p < 0.0001.

Analysis of fibre orientation in these images with OrientationJ^19^ and TWOMBLI^20^ revealed significantly higher local coherence in both OX and KO matrices compared to control (Fig. 1, C). The total area of collagen I signal, length of fibres and number of fibre endpoints were increased in KO samples which also demonstrated a decrease in lacunarity, confirming KO matrices are more friable and comprised of many individual dissociated fibrils (Fig. 1, C). Concurrently, OX samples exhibited a decrease in fibre branchpoints, corroborating the presence of large continuous fibres compared to WT (Fig. 1, C).

### Intermolecular crosslinks prime collagen molecular conformation and dynamics for cell-directed forces

To investigate the molecular basis for changes in collagen I microarchitecture we performed solid-state nuclear magnetic resonance (SSNMR) spectroscopy in fully hydrated *in vitro* matrices. To explore the molecular architecture of collagen I specifically we generated U-^13^C glycine (Gly) and L-proline (Pro) labelled samples as Gly, Pro and hydroxyproline (Hyp) have particularly high abundance in fibrillar collagens: Gly is every third amino acid residue (∼33%), and Pro is also widely distributed. In contrast, 40% of Hyp is in GPO triplets which our previous work has shown are arranged in collagen type I fibrils in bands in close proximity to integrin binding sites^21,22^.

^13^C{^1^H} cross polarization (CP) spectra demonstrated increased normalised signal intensity (when intensities are normalised to Hyp C_γ_ as a measure of collagen content) for OX and KO samples compared to WT at chemical shifts on the high frequency side of isotropic signal maxima assigned to Gly, Pro and Hyp carbons in the triple helix (Fig. 2, A; dashed lines).

**Figure 2:**
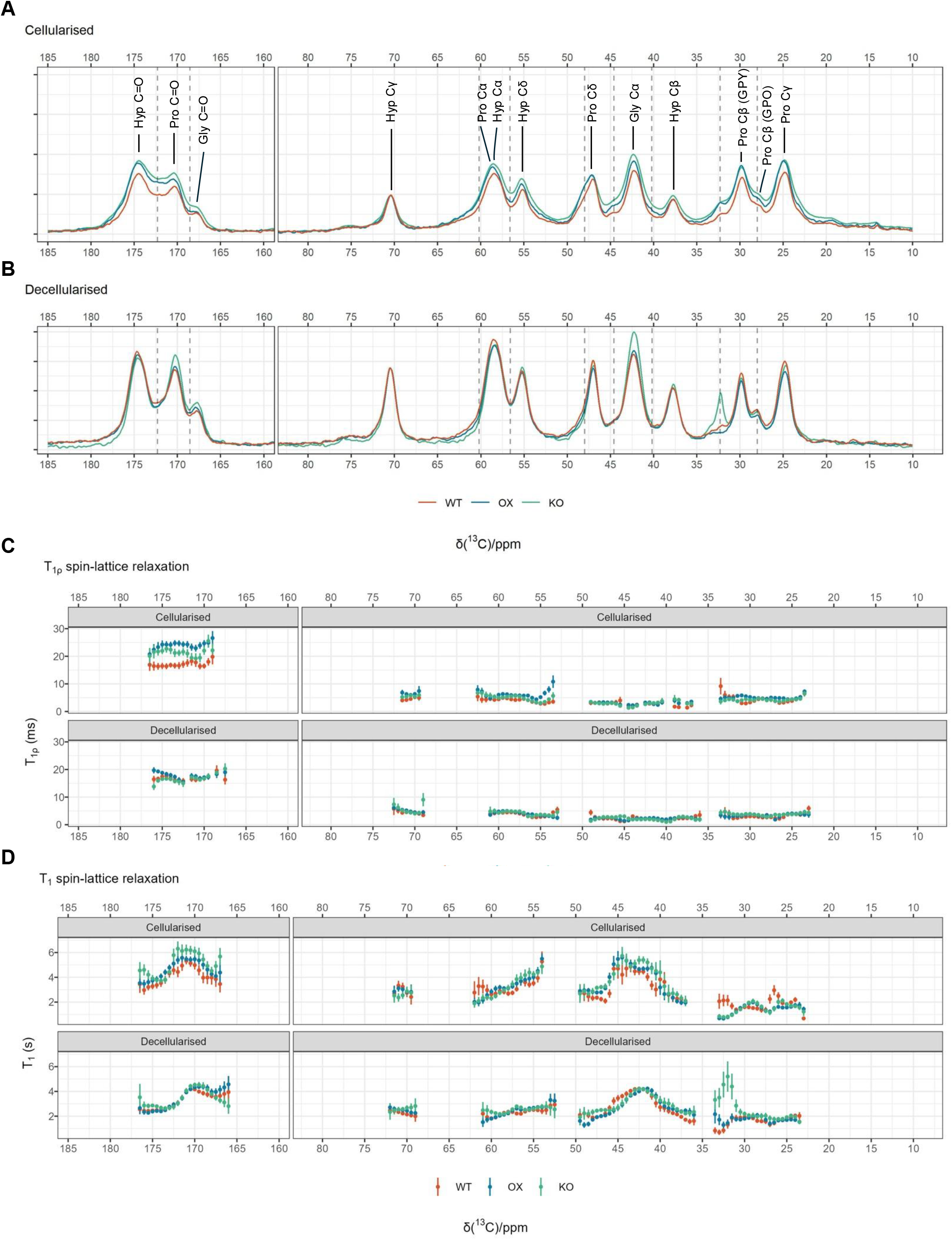
Altered intermolecular crosslinking and cell-mediated forces shape collagen I molecular conformation and dynamics. **(A,B)** _13_C{^1^H} cross polarization (CP) spectra in cellularized and decellularized U-^13^C glycine and L-proline labelled *in vitro* extracellular matrices. CP signal intensities are normalized to Hyp_γ_. **(B)** ^13^C T_1_ spin-lattice relaxation and **(C)** ^13^C T_1ρ_ spin-

Intensity differences were persistent when signals were normalised to the adjacent isotropic signal maximum (Supp. Fig. 2). Amino acid ^13^C chemical shifts are sensitive to protein secondary structure, with triple helices giving rise to lower shifts than other folding regimes for backbone and Cβ carbons^23–25^. Intensity at higher chemical shifts from the expected triple helix isotropic peaks therefore suggests regions of non-canonical collagen secondary structure in samples with increased and decreased crosslinking relative to WT.

Cells are known to actively apply traction forces to the ECM through adhesion complexes causing deformation of fibrous matrix components including collagen I^26–28^. We therefore performed decellularization of isotopically-labelled matrices to separate the effects of altered intermolecular crosslinking with cell-driven deformations to secondary structure through adhesion. Remarkably, decellularized ^13^C{^1^H} CP spectra demonstrated highly similar spectral chemical shifts and intensity patterns throughout the spectra between all samples, aside from small increases at Pro C’ and Gly C’ and C_α_, and a sharp peak at 32.3 ppm in KO samples (Fig. 2, A). We have previously shown that mechanical strain in collagen triple helices is predicted to induce shifts in backbone carbonyl carbon isotropic ^13^C chemical shifts consistent with the observed shifts in the pattern of carbonyl signal intensity in cellularised OX and KO samples^29^. These data suggest that crosslinking primes collagen I molecules into altered, non-canonical conformations detectable by SSNMR, and these conformational changes are further modulated by cell-driven traction forces from adhesion.

CP spectral intensities are sensitive to molecular dynamics on the μs timescale. When cellularised CP spectral intensities are normalised to Hyp C_γ_ (Fig. 2, A), Pro C_β_ or Pro C_y_ (Supp. Fig. 2, B-C), signals whose CP intensity are expected to be relatively unaffected by any μs timescale motion because of the fast Pro and Hyp ring flipping which dominates these sidechains, we find that the intensity of spectra for OX and KO samples are significantly higher across the whole spectrum compared to the spectra for WT samples. This suggests that there is more collagen molecular motion on the μs timescale for molecules in WT samples than KO or OX. To further probe this, we performed relaxation time measurements in cellularised and decellularized samples to investigate the contribution of collagen molecular dynamics to observed intensities. Milli - nanosecond-scale motions for collagen backbone carbonyl ^13^C were measured through ^13^C T_1ρ_ relaxation times under cross polarization (50kHz spin lock pulse) resolved by ^13^C chemical shift, revealing significantly faster relaxation for throughout the carbonyl spectral region for WT samples compared to OX and KO samples, suggesting faster dynamics or greater amplitude of dynamics on this timescale or both, in WT collagen fibrils (Fig. 2, B). We also investigated faster collagen molecular motions (nanoseconds – picoseconds) with ^13^C T_1_ spin-lattice relaxation, again resolved according to chemical shift which demonstrated a pronounced decrease in T_1_ relaxation times at non-triple helical Pro ^13^C sites in genetically-modified samples at 32.3 ppm (2.1-fold ± 0.20, Pro C_γ_) and 26.3 ppm (1.52-fold ± 0.31, Pro C_γ_), indicating increased nanosecond-scale molecular motion in LOX OX and KO collagen (Fig. 2, C). We find that in this faster timescale, there is also greater extent of collagen backbone carbonyl motion in WT compared to KO and OX samples.

Collectively, these data demonstrate that collagen intermolecular crosslinking and cell adhesion are functionally interlinked in regulating the range of accessible structural conformations and molecular dynamics of collagen I. Alterations in intermolecular crosslinking result in localized unfolding of the collagen triple helix, changes in the extent of triple helix compression and changes in both nanosecond- and microsecond-scale molecular motions.

However, these structural and dynamic changes are largely reversed upon decellularization, indicating that cell-mediated forces play a critical role in modulating collagen molecular architecture in conjunction with crosslinking. The persistence of a signal at 32.3 ppm from a particularly immobile component in KO samples following decellularization further suggests that certain conformational states may be stabilized by reduced crosslinking and that cell traction forces may be important in accessing these conformational states. These results highlight a mechanistic coupling between biochemical crosslinking and mechanical cell-matrix interactions that collectively define the conformational and dynamic landscape of collagen I.

### Altered collagen molecular structure and dynamics informs cell-matrix interactions and mechanosignalling

Having established that intermolecular crosslinking in collagen primes its range of molecular conformations and their dynamics, we then investigated how the extent of collagen intermolecular crosslinking affected cell adhesion using confocal microscopy.

Cytoskeletal staining revealed abundant F-actin stress fibres in OX cells (Fig. 1, A; white arrowheads), while KO cells exhibited accumulation of cortical actin (Fig. 3, A; white arrowheads). Immunofluorescence for the focal adhesion marker vinculin was used to quantify the number and morphology of adhesions. Modified cells demonstrated a significant reduction in the number of adhesions (p < 0.05) compared to WT controls. Adhesions were significantly larger in area (p < 0.001) and perimeter (p < 0.05) in OX cells compared to controls, while KO samples exhibited a decrease in adhesion area (p < 0.05) and perimeter (p < 0.001) (Fig. 3, B).

**Figure 3:**
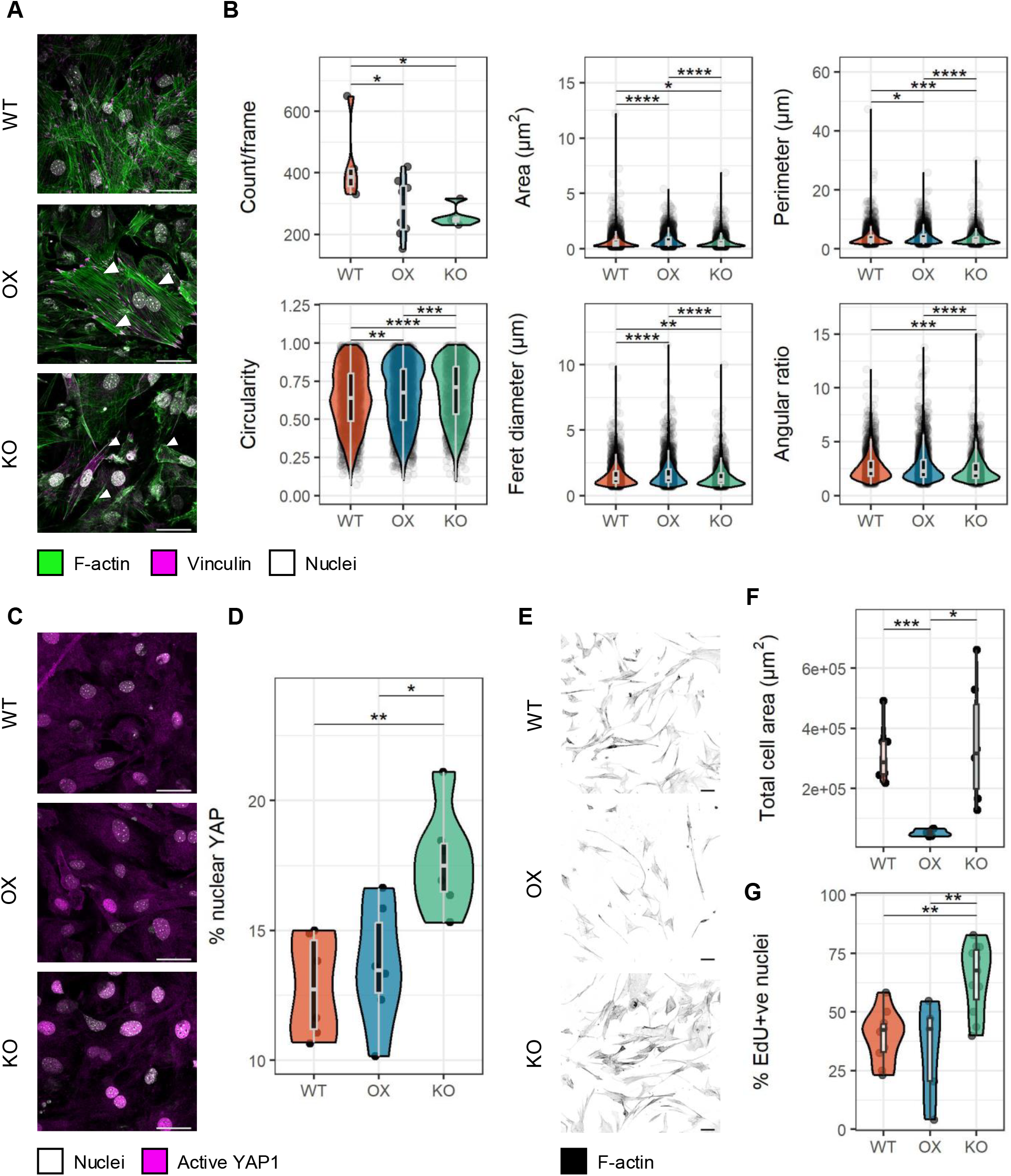
Intermolecular crosslinking in collagen regulates cell-matrix interactions through focal adhesions. **(A,B)** Immunoflouresence staining for the focal adhesion marker vinculin demonstrates alterations in the number and morphology of adhesions depending on the degree of intermolecular collagen crosslinking. **(C,D)** Immunoflouresence staining for active YAP1 shows an increase in its nuclear localisation in LOX KO cells. **(E,F)** WT naive cells replated on decellularized ECMs show a reduction in cell spreading on highly crosslinked (OX) collagen matrices. **(G)** Replated WT cells exhibit an increase in proliferation on undercrosslinked (KO) matrices. In imaging data n = 6 images/genotype (A-C) or 10 images/genotype (E,F) from at least 2 independent experiments. Statistical testing performed using pairwise Tukey Honest Significant Difference tests with Bonferonni correction: ^*^ p < 0.05; ^**^ p < 0.01; ^***^ p < 0.001; ^****^ p < 0.0001. Scale bars = 40µm (A,C); 100µm (E).

Adhesions also displayed altered morphology, with more circular adhesions in KO cells, evidenced by significant increases in circularity (p < 0.01) and reductions in angular ratio (p < 0.01) compared to controls (Fig. 3, B).

To analyse changes in cellular mechanosignalling as a result of altered adhesion, we quantified the nuclear localisation of active YAP1 as a proxy for inactivation of the mechanosensitive Hippo signalling pathway. Surprisingly, KO cells demonstrated a significant increase in active YAP1 nuclear localisation compared to WT (p < 0.01) and OX (p < 0.05) cells.

To disentangle the effects of cell-intrinsic effects during collagen fibrillogenesis and crosslinking, we decellularized matrices and reintroduced naïve WT cells to assess their ability to adhere and spread on ECM harbouring distinct levels of crosslinking. OX matrices significantly inhibited cell spreading after 48h of culture compared to WT controls (p < 0.001) or KO matrices (p < 0.05) (Fig. 3, E,F). While cells on KO matrices showed no difference in spreading compared to controls, they did exhibit an increase in EdU positive nuclei (p < 0.01), consistent with inhibition of Hippo signalling through increased active YAP1 nuclear localisation stimulating cell proliferation (Fig. 3, G).

These findings demonstrate that the degree of intermolecular collagen crosslinking acts as a potent regulator of cell–ECM interactions, reshaping cytoskeletal architecture, adhesion organisation, and mechanosignalling in a manner not solely explained by bulk stiffness.

Increased crosslinking (OX) promoted robust F-actin stress fibre formation and large, mature focal adhesions, yet paradoxically restricted cell spreading on decellularised matrices, consistent with steric or topographical constraints on integrin engagement. In contrast, reduced crosslinking (KO) yielded smaller, more circular adhesions and cortical actin enrichment, but unexpectedly drove elevated nuclear localisation of active YAP1 and increased proliferation, both in native and decellularised matrices, indicating Hippo pathway inactivation. Together, these results reveal that collagen crosslinking patterns encode complex mechanical and structural cues that differentially control adhesion maturation, cytoskeletal tension, and proliferative signalling, positioning ECM crosslinking as a tuneable instructive element in tissue mechanobiology.

## Discussion and conclusions

Our results reveal that LOX–mediated collagen crosslinking encodes structural and mechanical instructions that extend beyond matrix stiffening. By genetically modulating LOX in collagen I– producing cells, we show that crosslink density dictates fibrillar architecture, accessible molecular conformations and dynamics, and downstream mechanosignalling in a mechanically-tuned biochemical code.

Enhanced crosslinking (OX) produced thick, continuous bundles of aligned fibrils, while loss of crosslinking (KO) yielded friable, dissociated fibrils. These structural signatures match prior reports that LOX-derived covalent bonds stabilise fibril fusion and tensile strength^30^ and that their absence compromises fibre assembly^31–33^. Unexpectedly, both OX and KO matrices exhibited increased local fibril orientation, suggesting that altered crosslinking can sharpen local alignment through cell-driven tension during deposition^34^.

SSNMR uncovered local triple-helix unfolding and triple helix compression in both OX and KO matrices, with altered nanosecond- and microsecond-scale molecular motions that were largely reversed upon decellularisation, implicating a synergy between crosslinking chemistry and cell-applied forces. The persistence of a Pro_β_ signature from highly immobile, likely unfolded collagen in KO matrices after decellularisation suggests that reduced crosslinking stabilises certain unfolded states independently of active loading. We have previously shown that fast molecular motions at proline rings provide fibril flexibility through bands of GPO triplets organised across the fibril structure around integrin binding sites^21,22^ and that only relatively small distortions in triple helix longitudinal alignment are required to annul stabilising charge-charge interactions between individual tropocollagen molecules^35^. The data presented here therefore suggest that LOX-mediated intermolecular crosslinking organises the topology of mechanically sensitive domains within the fibril. By dictating where local triple-helix unfolding can occur under load and the dynamics of available binding sites, crosslinking chemistry may tune the accessibility and mechanical coupling of adhesion-relevant motifs, thereby pre-configuring the matrix for specific mechanochemical feedback loops during cell–ECM interactions.

These molecular changes translate into distinct cell–matrix behaviours. OX cells formed large, mature focal adhesions and robust stress fibres, yet naïve cells spread poorly on OX decellularised matrices – indicating steric, topographical or dynamic restrictions to integrin engagement^36^. KO cells, in contrast, formed small, circular adhesions but exhibited elevated nuclear YAP1 and proliferation, defying the canonical link between adhesion size and Hippo activation^37^. The influence of mechanical force sensed by intracellular components in the initiation and maturation of cell-matrix adhesions is well established^38,39^, however these data suggest that collagen dynamic molecular conformation may be a critical component in translating the structural and mechanical state of the ECM.

Collectively, these findings position collagen crosslinking as a tuneable regulator of the ECM’s “mechanical code”, integrating collagen molecular conformation and dynamics to direct cell adhesion.

## Acknowledgements

The study was funded by the European Union (ERC, EXTREME 101019499). Views and opinions expressed are however those of the author(s) only and do not necessarily reflect those of the European Union or the European Research Council Executive Agency. Neither the European Union nor the granting authority can be held responsible for them.

## Supplementary information

## Materials & methods

### Materials

Triton X-100, ammonium hydroxide, Tris-HCl, MgCl_2_, CaCl_2_, paraformaldehyde, bovine serum albumin (BSA), Tween-20, CuSO_4_ and hexamethyldisilazane (HMDS) were purchased from Sigma.

### Tissue culture

Reagents for tissue culture were purchased from Gibco unless otherwise indicated.

Mouse pre-osteoblast MC3T3-E1 cells (ATCC) were used to generate *in vitro* collagenous extracellular matrix (ECM). Cells were cultured at 37^°^C in a humidified 5% CO_2_ atmosphere in complete growth media comprising α-Minimum Essential Media (αMEM) with nucleosides supplemented with 10% heat-inactivated foetal bovine serum (FBS; PAN Biotech), 5 units/ml penicillin, 1µg/ml streptomycin and 0.292 mg/ml L-glutamine. Growth media was refreshed every 2-3 days.

To induce matrix production growth media was supplemented at cell confluence with 50 µg/ml L-ascorbic acid and matrices grown for 21 days. For ^13^C isotopic enrichment of *in vitro* ECM, osteogenic media was further supplemented with 50μg/ml U-^13^C glycine and 40μg/ml U-^13^C L-proline (Cambridge Isotope Laboratories) for the entire post-confluence culture period.

### Generation of lysyl oxidase knock-out and overexpression MC3T3s

CRISPR-Cas9 gene editing was used to generate *Lox* knock-out MC3T3s. Guide RNAs (gRNAs) were designed against *Lox* exon 1 and conjugated with trans-activating CRISPR RNAs (tracrRNAs) and recombinant Cas9 nuclease to form functional ribonucleoprotein complexes according to manufacturer’s instructions (Integrated DNA Technologies). A pool of 3 gRNAs was used for transfections. Complexes were reverse transfected using Lipofectamine CRISPRMAX (Thermo Fisher Scientific) according to manufacturer’s instructions and single clones generated by limiting dilution in 10% conditioned growth media. DNA was extracted form single clones using DNeasy spin columns (Qiagen) and sequenced using Sanger sequencing to validate successful edits.

The PiggyBac transposon system was used to generate MC3T3s overexpressing LOX. A donor plasmid encoding full-length mouse *Lox* upstream of a CMV promotor and blasticidin resistance upstream of a CMV promotor flanked by 3’ and 5’ inverted terminal repeat (ITR) sequences, and a helper plasmid encoding hyperactive PiggyBac transposase were purchased as *E*.*coli* glycerol stocks (Vector Builder). Plasmids were purified from overnight inoculations using Miniprep spin columns (Qiagen) and cells reverse transfected using Lipofectamine 3000 (Thermo Fisher Scientific) according to manufacturer’s instructions. Successfully transformed cells were selected by supplementing media with 10µg/ml blasticidin S (Santra Cruz Biotechnology) for 10 days and single clones generated by limiting dilution in 10% conditioned growth media.

### Quantitative reverse transcription polymerase chain reaction

RNA was isolated from cells at the end of the culture period using Trizol (Thermo Fisher Scientific) extraction followed by purification with RNeasy spin columns (Qiagen) according to manufacturer’s instructions. Reverse transcription was performed using the GoScript reverse transcriptase system (Promega) and quantitative polymerase chain reaction (qPCR) carried out with PowerTrack SYBR Green master mix (Thermo Fisher Scientific) using a CFX96 real-time thermal cycler (BioRad). The following primers were used for qPCR: *Actb*, forward: 5’-TGACGTTGACATCCGTAAAG-3’, reverse: 5’-GAGGAGCAATGATCTTGATCT-3’; *Gapdh*, forward: 5’-TGACCTCAACTACATGGTCTACA-3’, reverse: 5’-CTTCCCATTCTCGGCCTTG-3’; *Lox*, forward: 5’-AGTAATCTGAGGCCACCCAG-3’, reverse: 5’-AATAGGGGTTGTCGTCGGAG-3’. Data were analysed according to the ΔΔCt method using *Actb* and *Gapdh* as housekeeping controls.

### In vitro matrix decellularization

To decellularize *in vitro* matrices cells were washed in Dulbecco’s phosphate buffered saline (DPBS; Gibco) and frozen at -20^°^ C overnight. Samples were thawed at room temperature for 30min and treated with 0.5% Triton X-100; 20mM ammonium hydroxide in DPBS for 10min at 37^°^C. Detergent solution was diluted 1:2 in DPBS and samples incubated overnight at 4^°^C. Samples were washed well in DBPS and treated with 10 µg/ml DNAseI (Sigma) in 100mM Tris-HCl; 25mM MgCl_2_; 1mM CaCl_2_ pH 7.6 for 30min at 37^°^C. Samples were stored at -20^°^C before downstream analysis.

To prepare decellularized *in vitro* matrices for cell replating, culture surfaces were gelatine coated prior to tissue culture to prevent matrices lifting after detergent treatment as previously described^40^. Briefly, 0.2% gelatine (Sigma) in DPBS was dissolved by autoclaving and filtered using a 0.22 µm syringe filter (Sigma). Cell culture plastic or glass coverslips were incubated in gelatine solution for 1 hour at room temperature, washed in DPBS and gelatine crosslinked in 1% glutaraldehyde in DPBS for 30min. Glutaraldehyde was quenched in 1M ethanolamine in DPBS for 30min and surfaces washed well in DPBS and stored at 4^°^C before use.

For replating experiments, decellularized matrices were prepared as above in 12-well tissue culture plates and WT cells replated in growth media at a density of 10,000 cells/well. Cells were cultured for 48h and prepared as appropriate for downstream analysis.

### Immunofluorescence

For immunofluorescence cells were fixed in 4% paraformaldehyde in DPBS for 10min at room temperature, washed well in DPBS and stored at 4^°^C before staining. Samples were blocked in 5% BSA in DPBS for 1 hour at room temperature and permeabilised in 0.5% Triton X-100 in DPBS for 30min at room temperature. Samples wee incubated with primary antibodies against collagen I (1:200; Abcam; EPR24331-53), vinculin (1:500; Thermo Fisher Scientific; 42H89L44), or active (non-phosphorylated) Yes1 associated transcriptional regulator (YAP1) (1:500; Abcam; ab205270) in 1% BSA; 0.05% Tween-20 for 48 hours at 4^°^ C in a humidity chamber. Primary antibodies were detected using an anti-rabbit IgG secondary antibody conjugated to AlexaFluor 647 (1:500; Abcam; ab150083) in 0.05% Tween-20 for 1 hour at room temperature. To visualise the actin cytoskeleton, samples were incubated with 100nM Acti-stain phalloidin (Universal Biologicals) simultaneous with the secondary antibody. Samples were mounted on microscope slides in Fluoroshield mounting medium with DAPI (Sigma) and stored at 4^°^C in darkness before imaging.

For EdU staining in replated samples, growth media was additionally supplemented with 10µM EdU (Abcam) for 48h. Cells on coverslips were fixed as above and EdU detected using a click chemistry reaction in a solution containing 2mM CuSO_4_, 8µM FAM-azide (Abcam) and 200mg/ml L-ascorbic acid for 30min at room temperature in darkness. Coverslips were mounted as above.

Laser scanning confocal microscopy was performed using a Stellaris 5 (Leica) microscope equipped with 20X air- and 40X- and 63X oil-immersion objective lenses.

### Scanning electron microscopy

Decellularized matrices on glass coverslips were fixed in 4% paraformaldehyde; 2.5% glutaraldehyde pH 7.6 in DPBS for 10min at room temperature, fixative refreshed and samples stored at 4^°^C. Samples were dehydrated through an ascending ethanol series and subsequently incubated in HMDS twice for 30min at room temperature, before the solvent was removed and samples left to air dry overnight.

Dried matrices were mounted onto aluminium sample stubs with adhesive carbon tabs (Agar Scientific) and electrical connection established with the sample surface using copper tape (Electron Microscopy Sciences). The sample surface was coated with 10nm platinum using a Q150T ES carbon coater (Quorum Technologies).

Samples were imaged using a CLARA2 (TESCAN) field emission gun scanning electron microscope (SEM) equipped with Everhart-Thornley and in-lens secondary electron detectors and operating at 5kV at a sample working distance of 5mm.

### Image analysis

OrientationJ^19^ for Fiji^41^ was used for local coherence analysis of collagen I immunofluorescence images. To generate orientation and vector field maps a σ of 2px with a cubic spline gradient was used in a 50×50 grid. Grid squares with a measured energy < 0.1 were discarded to remove background regions with no collagen I signal. TWOMBLI^20^ was used to quantify collagen fibril branching parameters with a line width of 10px, minimum branch length of 10px and a curvature window of 40px.

A custom Fiji macro was written to quantify focal adhesion number and morphology. Briefly, the vinculin fluorescence signal was subject to background subtraction, local contrast enhancement (blocksize = 19px; histogram bins = 256; maximum slope = 6), an exponential filter, and a median filter (σ = 2px). Automatic thresholding was performed using the Otsu algorithm and masked binary particles analysed.

Nuclear localisation of active YAP1 was quantified as a percentage of integrated fluorescence intensity within thresholded DAPI-stained nuclear boundaries to total integrated fluorescence intensity. Cell spreading was quantified as total thresholded area occupied by cells after a median filter (σ = 20px).

### Solid state nuclear magnetic resonance spectroscopy

Consumables for solid state nuclear magnetic resonance (SSNMR) spectroscopy were purchased from Bruker.

*In vitro* matrix samples were centrifuged at 300 × g for 3min at room temperature and bulk water removed. Fully hydrated matrices were packed into 3.2mm ZrO_2_ NMR rotors and capped with a VESPEL drive cap. Rotors were snap frozen on dry ice and stored at -20°C prior to analysis.

SSNMR was performed using an AVANCE Neo 400MHz (9.4T) ^1^H Larmor frequency wide bore spectrometer (Bruker) equipped with a 3.2mm Efree magic angle spinning (MAS) probe operating at 10kHz at a corrected sample temperature of -5^°^C. Sample temperature under MAS was corrected using the temperature-dependent T_1_ relaxation of KBr at the same conditions. ^13^C chemical shifts were referenced to crystalline α-glycine C_α_ at 43.1ppm at -5^°^C, relative to tetramethylsilane (TMS) at 0ppm.

In all experiments ^1^H 90^°^ pulses were performed at 100kHz and data were acquired with *SPINAL64* ^1^H heteronuclear decoupling at 100kHz. The recycle delay in all experiments was 2.5s. ^13^C{^1^H} cross polarisation (CP) spectra were acquired with a *tanhc60* shaped ^1^H pulse and a square ^13^C pulse optimised for total signal intensity at 98.5kHz with a contact time of 2ms, acquisition time of 30ms and 3072 scans. ^13^C T_1_ spin-lattice relaxation was quantified using a CP inversion recovery approach using the same CP parameters and a total of 24 T_1_ delays between 0.1s-100s and 256 scans. ^13^C T_1ρ_ spin-lattice relaxation under a CP spin lock was quantified using a CP spin lock approach using the above CP parameters and a 50kHz ^13^C spin lock field with a total of 24 T_1ρ_ delays between 1ms-100ms and 512 scans.

Relaxation data were deconvoluted according to ^13^C chemical shift. Normalised signal intensity was integrated at every 0.5ppm between 10-190ppm for each relaxation delay and the relaxation constant fitted using a nonlinear least squares approach according to the functions:

T_1_ spin-lattice relaxation:

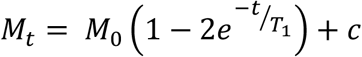

Where *M*_*t*_ is longitudinal magnetization at delay time *t* (proportional to integrated signal intensity), *M*_*0*_ is the equilibrium longitudinal magnetization, *T*_*1*_ is the spin-lattice relaxation constant and *c* is a constant to correct for noise.

T_1ρ_ spin-lattice relaxation under spin lock:

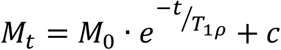

Where *T*_*1ρ*_ is the spin-lattice relaxation under spin lock constant and the other parameters are as above. Relaxation data were plotted as fitted estimate ± standard error of the fit according to ^13^C chemical shift. NMR data processing and fitting was performed in R.

**Supplementary Figure 1:**
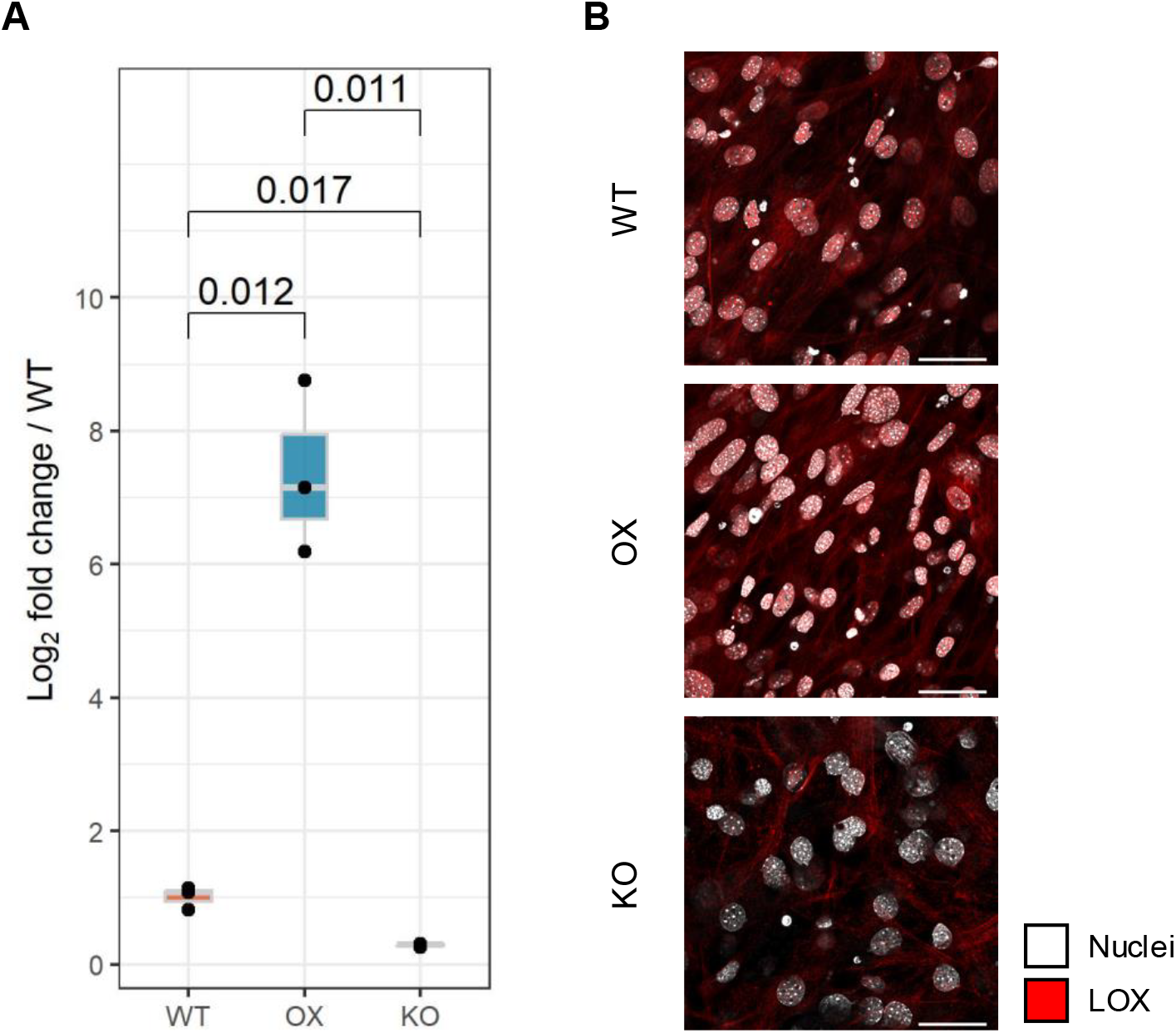
Validation of genetic manipulation of *Lox* expression in MC3T3 cells. **(A)** Quantitative reverse transcription polymerase chain reaction (RT-qPCR) for *Lox* mRNA in wild-type (WT), overexpression (OX) and knock-out (KO) cells at 21 days post-confluence L-ascorbic acid treatment. Data are expressed as Log_2_ fold change compared to average WT values. Statistical testing performed using a t-test. **(B)** Immunofluorescence staining for LOX. Scale bars = 40µm.

**Supplementary Figure 2:**
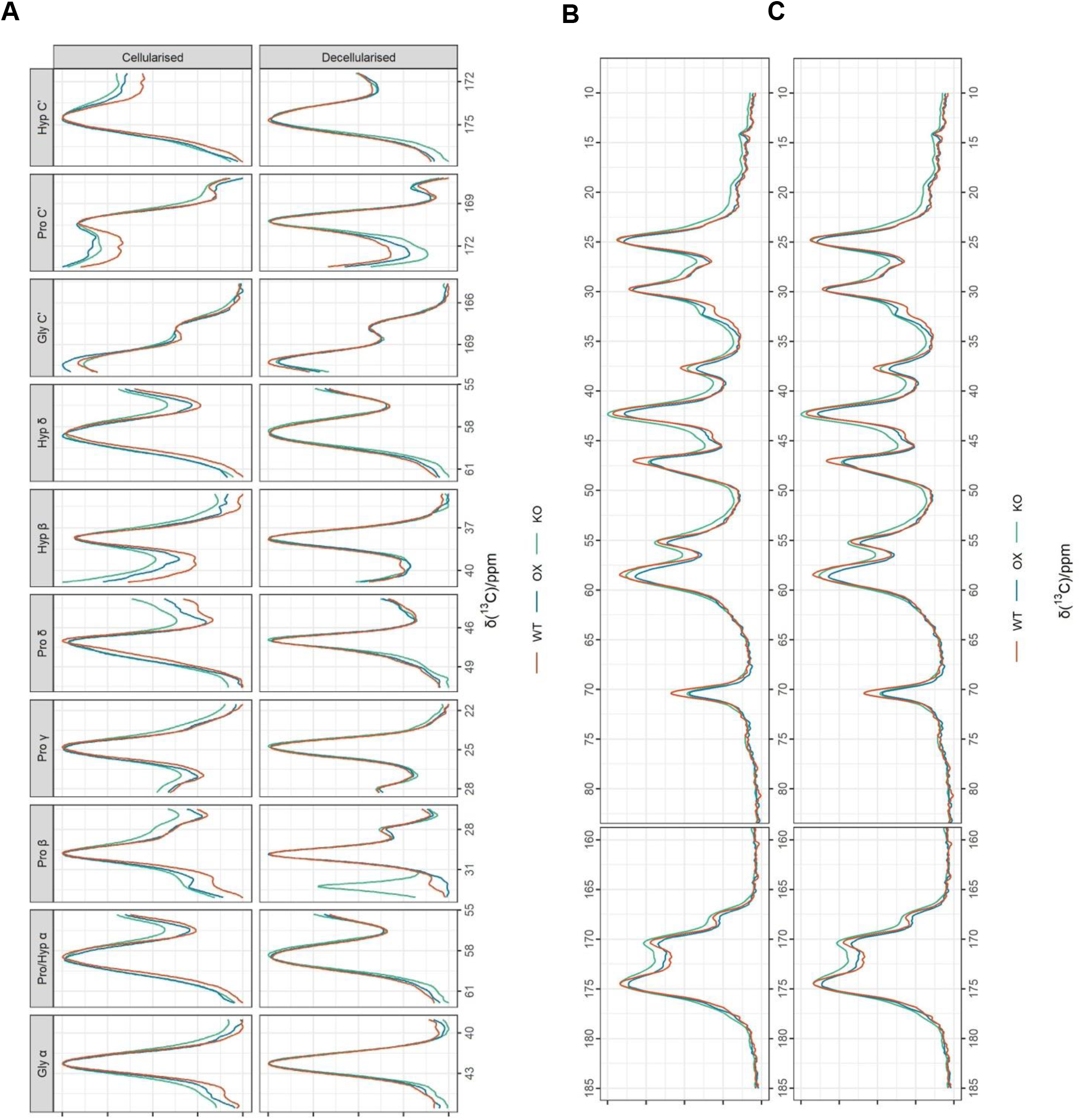
Solid state nuclear magnetic resonance (SSNMR) spectroscopy of *in vitro* MC3T3 hydrated matrices. **(A)** ^13^C{^1^H} cross polarisation (CP) spectra of hydrated MC3T3 U-^13^C labelled glycine and L-proline *in vitro* extracellular matrices as in Fig. 2, A-B. Signal intensities are normalised to the isotropic peak shown in each panel. **(B,C)** Full ^**13**^C{^1^H} CP spectra as above with signal intensities normalised to Pro C_β_ (B) and Pro C_γ_ (C).

